# Real-Time Quantitative PCR Method for Assessing Wheat Cultivars for Resistance to *Zymoseptoria tritici*

**DOI:** 10.1101/2025.04.02.646880

**Authors:** Tika B. Adhikari, Boovaraghan Balaji, Stephen B. Goodwin

## Abstract

Septoria tritici blotch, caused by *Zymoseptoria tritici* (formerly *Mycosphaerella graminicola*), is an economically significant disease of wheat (*Triticum aestivum*) worldwide. However, there is little understanding of the fungus’ growth dynamics during the 14- to 18-day latent period between penetration and symptom expression, making it challenging to develop wheat cultivars resistant to *Z. tritici* Furthermore, environmental factors and variations in disease-scoring systems among evaluators add to the complexity. To address these issues and quantify the fungus’ growth during the initial stages of infection, we developed a real-time quantitative polymerase chain reaction (real-time PCR) method to monitor the *T. aestivum - Z. tritici* pathosystem. The assay used specific primers designed from ß-tubulin gene sequences of *Z. tritici* to quantify fungal DNA in susceptible and resistant wheat cultivars and segregating recombinant-inbred lines (RILs) that were inoculated at seedling and adult-plant stages with low or high concentrations of inoculum. The real-time PCR method was compared with visual disease assessment for 0 to 27 days after inoculation (DAI). The results showed that fungal DNA increased more quickly in two susceptible cultivars than in resistant cultivars with the *Stb4* or *Stb8* genes for resistance. In the susceptible cultivars, the amount of fungal DNA remained low until symptoms became visible at around 18 DAI. Disease severity and fungal DNA in the two resistant cultivars were less than in either susceptible cultivar, starting at 12 DAI. The differences in fungal DNA between resistant and susceptible cultivars were more significant in adult plant tests that used a higher concentration of inoculum. The data analyses showed that the fungus was not eliminated during resistant interactions but could persist throughout the 27 days. Our results suggest that the real-time PCR method can distinguish between resistant and susceptible cultivars starting at 12 DAI and can be used to evaluate early-stage breeding materials for both quantitative and qualitative resistance to *Z. tritici*.

## Introduction

Septoria tritici blotch (STB), caused by *Zymoseptoria tritici* (formerly *Mycosphaerella graminicola*; anamorph *Septoria tritici*), is a widespread and economically significant disease of wheat (Eyal et al., 1985; 1987). STB is a serious threat to wheat production in the USA, particularly in the Pacific Northwest (Jackson et al., 2000) where the environment is favorable for disease development almost every year, and in other growing regions when spring rains are favorable for disease development. Wheat cultivars resistant to *Z. tritici* restrict or delay pathogen development (Nelson and Marshall, 1990). The disease can be spread by air-borne ascospores or splash-dispersed conidia, infecting the flag leaf during the wheat-growing season (Eyal et al., 1987; Ponomarenko et al., 2011). In disease-conducive environments, epidemics can quickly cause significant reductions in yield on susceptible plants (Gilbert et al., 1998; Shaner and Finney, 1976). Histopathological analysis has revealed that *Z. tritici* infects its host plant through stomata. During infection, the hyphae strictly remain in the intercellular space in close contact with mesophyll cells (Cohen and Eyal, 1993; Duncan and Howard, 2000; Kema et al., 1996). However, no specialized feeding structures like haustoria are formed. The first visible symptoms of the infection appear as chlorotic areas on the leaf surface about 10 days after inoculation (DAI) (Ponomarenko et al., 2011). These chlorotic areas later become necrotic lesions starting around 14 to 18 DAI. When the first signs of pycnidia appear, these structures emerge from the stomata and become filled with basket-shaped masses of mycelia (Cohen and Eyal, 1993; Duncan and Howard, 2000; Kema et al., 1996).

Wheat cultivars are usually assessed for resistance to *Z. tritici* by visual estimation of symptoms. However, this method has several disadvantages. First, the fungus has a long latent period (up to 14 or 18 days), so the standard time for evaluating resistance to *Z. tritici* in wheat cultivars is between 24 and 28 DAI. Although scoring adult plants’ flag leaves usually produces the best results (Adhikari et al. 2003), growing plants to maturity slows evaluation and limits the number of plants tested. Second, using an integer scale for rating resistance may vary from person to person, making it challenging to distinguish resistant plants phenotypically. Finally, the expression of resistance to *Z. tritici* under greenhouse and field conditions can be significantly influenced by the environment. Some of these difficulties can be minimized by detached-leaf assays (Arraiano et al. 2001), by scoring plants in a greenhouse (Kema et al. 1996) or by automating the phenotyping step (Stewart et al., 2016), but these require much more effort and reduced throughput. Therefore, a rapid, sensitive, and high-throughput method for quantifying fungal DNA would be helpful in accurately assessing pathogenicity and for large-scale testing of plant genetic resources for resistance to *Z. tritici*.

Several methods have been used for detecting and quantifying *Z. tritici* in susceptible and resistant wheat cultivars. Cohen and Eyal (1993) estimated mycelial growth histologically in the intercellular spaces of mesophyll cells of resistant and susceptible wheat cultivars. Although fungal growth increased dramatically in the susceptible response during the later stages of pathogenesis, no differences were observed at an early stage of the infection (8 DAI) (Kema et al. 1996). The histopathological analyses also showed that a non-virulent mutant colonized sub-stomatal cavities less efficiently and displayed reduced intercellular growth and concomitantly less fungal growth in the apoplast of wheat leaves compared to untransformed controls (Stergiopoulos et al. 2003). In another approach, ß-glucuronidase-producing transformants of *Z. tritici* were used to examine fungal development in inoculated wheat leaves. In general, the rates of fungal growth and the estimated levels of fungal protein when the pycnidia matured appeared to be high in compatible interactions and moderate to low in incompatible interactions (Pnini-Cohen et al. 2000).

A multiplex polymerase chain reaction (PCR) method was used to detect and quantify four foliar fungal pathogens of wheat, including *Z. tritici* (Fraaije et al. 2001; Guo et al. 2006). However, those primer pairs for each foliar pathogen varied in sensitivity due to weak non-specific amplification and primer dimer formation during the PCR method. Although some of the methods reported previously are promising, they are not sensitive enough to monitor the slight changes in fungal DNA that may occur during pathogenesis. The real-time PCR (Heid et al. 1996) method could be rapid, sensitive, and accurate enough to estimate the quantity of *Z. tritici* DNA in large numbers of inoculated wheat plants. Real-time PCR has been used successfully for the detection and quantification of a range of plant pathogens, such as bacteria (Brouver et al. 2003; Humphris et al. 2015; Peňázová, et al. 2020; Schaad et al. 1999; Weller et al. 2000), fungi (Bates et al. 2001; Brouwer et al. 2003; Chu et al. 2019; Gachon and Saindrenan 2004; Milgate et al. 2023; Oliver et al. 2008; Ozdemir et al. 2020; Qi and Yang 2002; Vandemark et al. 2002; Winron et al. 2003) and viruses (Balaji et al. 2003; Diaz-Lara et al. 2020). It has been used to estimate fungicide resistance in field populations of *Z. tritici* (Fraaije et al. 2005), to detect presymptomatic fungal biomass to guide fungicide applications (Guo et al. 2006; Tonti et al 2019) and to identify early responses of candidate genes in resistant cultivars (Farsad et al 2013).

Breeding wheat for resistance to *Z. tritici* could benefit from a comprehensive analysis of fungal DNA accumulation in susceptible (compatible) and resistant (incompatible) interactions. The ultimate goal of this study was to use real-time PCR as a disease assessment method to differentiate between resistant and susceptible wheat cultivars and to enhance our understanding of STB development. We hypothesized that the quantity of fungal DNA increases rapidly after infection of susceptible wheat cultivars by *Z. tritici*, but it should remain constant or increase only slightly in resistant cultivars. This change should be noticeable in the amount of fungal DNA present in total nucleic acids extracted from inoculated wheat leaves, potentially much earlier than symptom development. The specific objectives of this study were to: (i) develop an improved real-time PCR method for quantifying *Z. tritici* in infected wheat leaves; (ii) use the real-time PCR method to monitor fungal DNA accumulation over a 27-day time-course in two susceptible and two resistant wheat cultivars with different specific resistance genes; (iii) apply the real-time PCR method to examine the effect of inoculum concentration and plant age on fungal DNA accumulation; and (iv) determine the relationship between fungal DNA and disease severity in a population of segregating resistant and susceptible recombinant-inbred lines (RILs) to compare the real-time PCR quantification with visual disease-assessment methods.

## Materials and methods

### Selection of wheat cultivars

Three sets of wheat cultivars or populations were evaluated using a real-time PCR method. Set 1 consisted of two resistant (Tadinia and W7984) and two susceptible (Yecora Rojo and Opata 85) cultivars (Adhikari et al. 2003; Adhikari et al. 2004a). Tadinia is a spring wheat with the *Stb4* gene for resistance to *Z. tritici* (Adhikari et al. 2004a). The synthetic wheat W7984 has the gene *Stb8* (Adhikari et al. 2003). This standard set was used to monitor the DNA quantity of *Z. tritici* in the greenhouse over time. In addition, this set of cultivars was used in a separate experiment to determine the effect of low and high inoculum concentrations on fungal DNA. Set 2 consisted of 20 recombinant-inbred lines (RILs) from a three-way cross between the resistant cultivar Tadinia and the susceptible parent (Yecora Rojo × UC554). Ten resistant (pycnidial density scores < 1.0) lines and 10 susceptible (pycnidial density scores > 3.0) lines were selected from our previous analyses (Adhikari et al. 2004a). The utility of a real-time PCR method for quantifying fungal DNA was compared with visual disease assessment results (Adhikari et al. 2004a) to discriminate between resistant and susceptible parents and RILs. The third set consisted of two highly susceptible (Chinese Spring and Taichung 29) and two resistant (Tadinia and Veranopolis) cultivars. Chinese Spring contains the recently cloned *Stb6* resistance gene (Saintenac et al. 2018), but virulence to this gene is almost universally present among contemporary isolates of *Z. tritici* (Stephens et al. 2021) and this cultivar was highly susceptible to our Indiana tester isolate. Veranopolis possesses gene *Stb2* (Wilson 1979; 1985), which confers resistance to both Indiana and Australian isolates of *Z. tritici* (Adhikari et al. 2004c; Goodwin et al. 2015; Liu et al. 2013). Total fungal DNA detected in each cultivar at 6, 9 and 12 DAI was compared at seedling and adult-plant stages.

### Fungal inoculum preparation and plant inoculations

Plants were grown separately in a greenhouse for each experiment. All conditions and cultural practices were as described previously (Adhikari et al. 2003). The tester isolate IN95-Lafayette-1196-WW-1-4 of *Z. tritici* (also known as T48) was collected in 1995 from Lafayette, Indiana, USA, and was grown and maintained as a lyophilized filter paper stock, except that it was re-isolated from infected leaves occasionally to maintain pathogenicity. Spores prepared as inoculum were harvested by filtration through sterilized Whatman paper and transferred into sterile 1.5-ml microcentrifuge tubes to estimate a standard curve for fungal DNA. The spores were lyophilized and stored at -80° C before DNA extraction.

Inoculum was prepared by placing a 1-cm^2^ piece of agar culture into 250-ml Erlenmeyer flasks containing 100 ml of yeast-sucrose liquid medium (10 g each of sucrose and yeast extract per liter of distilled water) and 100 µl of kanamycin sulfate solution (25 mg/ml). Flasks were plugged with cotton and placed on an orbital shaker (Barnstead/Thermolyne, Dubuque, IA) at 150 rpm at room temperature for 3 to 5 days. The inoculum suspension for most experiments was adjusted to 3 to 6 × 10^6^ spores/ml by hemacytometer before inoculation. Plants were spray inoculated approximately seven weeks after sowing.

### DNA extraction

Fungal mycelia, pycnidiospores, or wheat leaves were ground with liquid nitrogen using a mortar and pestle. Approximately 100 mg of powder were transferred into 1.5-ml microcentrifuge tubes, and genomic DNA was extracted with the DNeasy Mini Kit (Qiagen, Valencia, CA) and quantified with a fluorometer (Hoefer Scientific Instruments, San Francisco, CA). DNA of major fungal pathogens of wheat and unrelated rice pathogens were obtained and included as negative controls (Table 1). Genomic DNA from pathogen-infected and healthy (uninoculated) wheat leaves was also extracted and included as positive and negative controls, respectively.

**TABLE 1.**
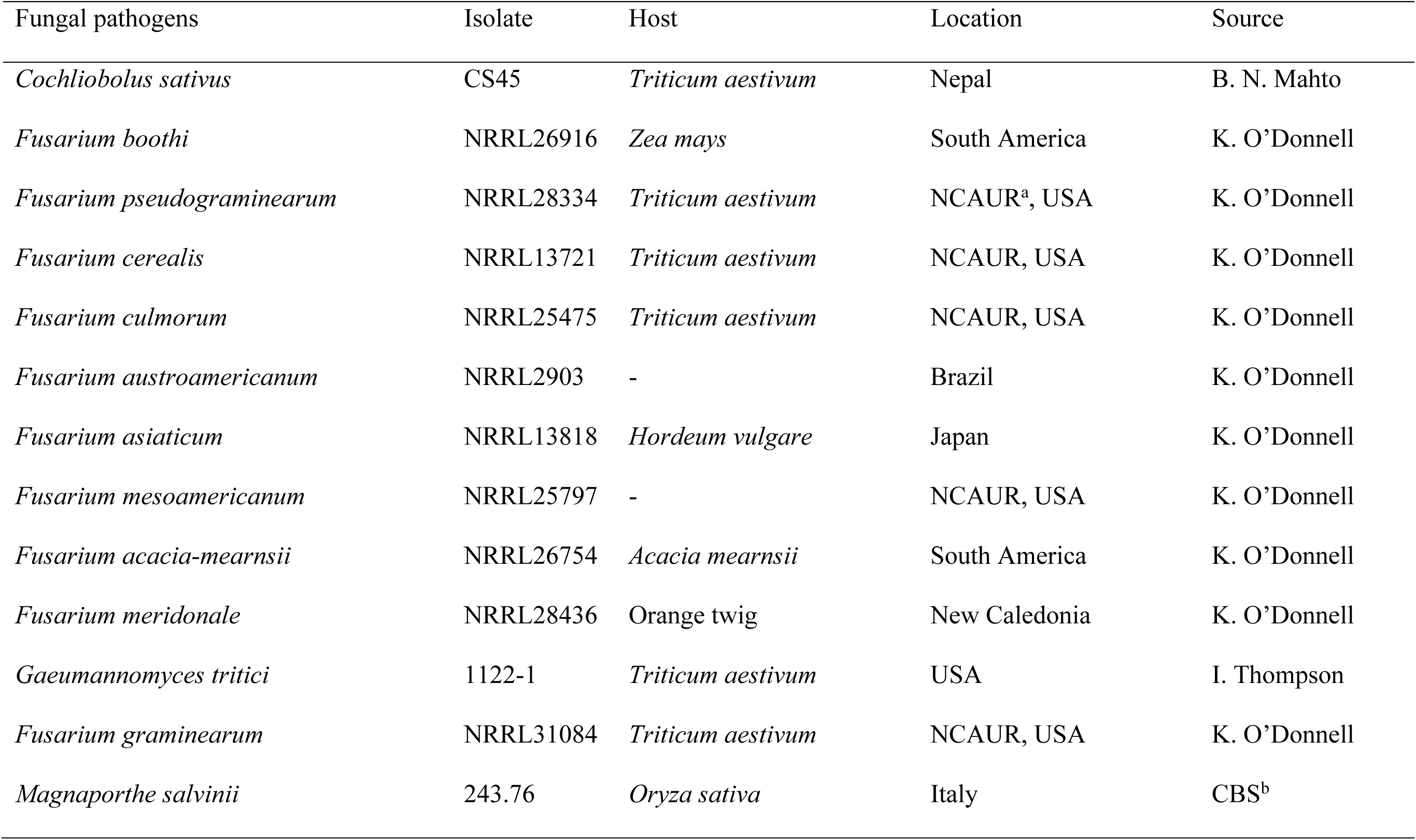
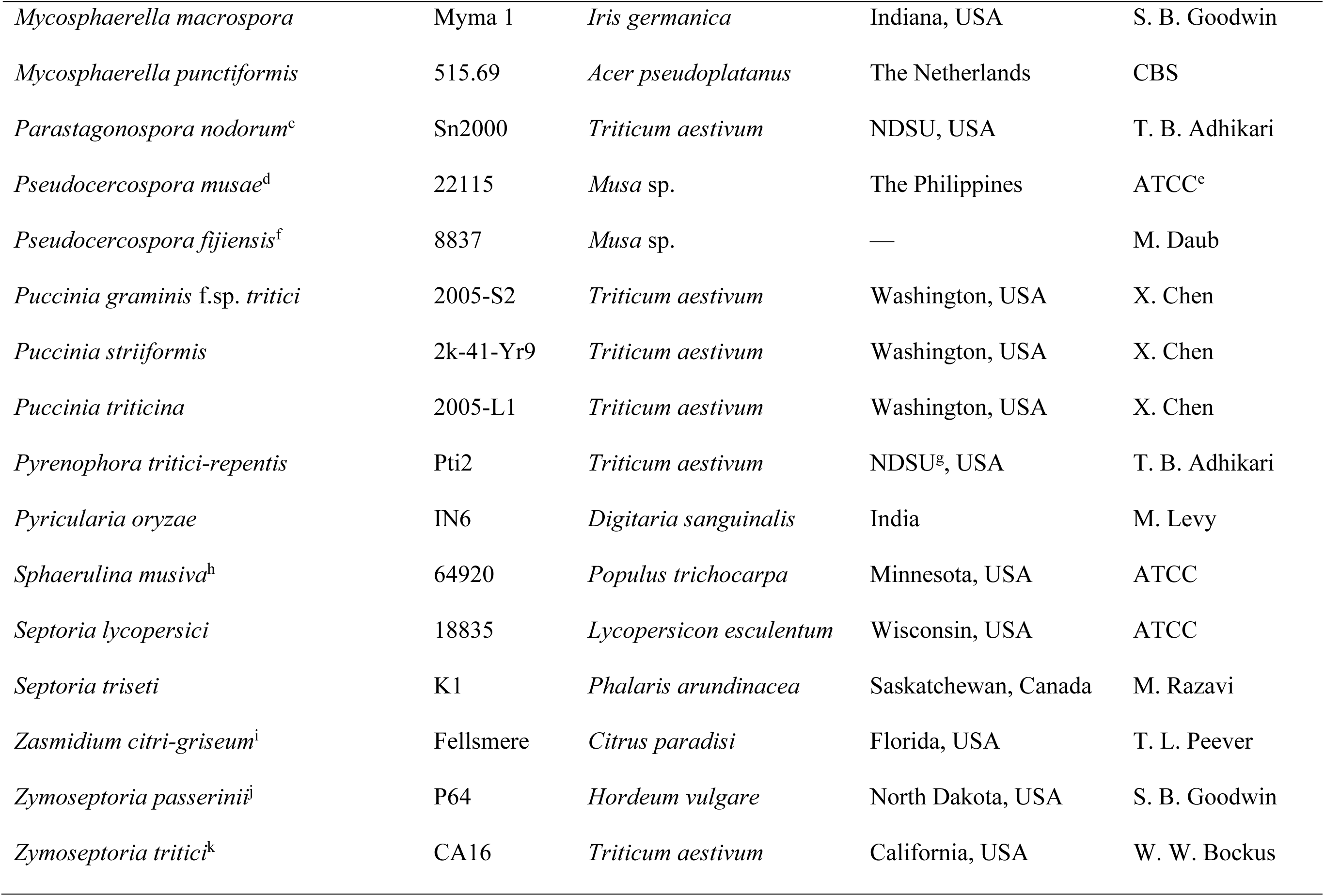
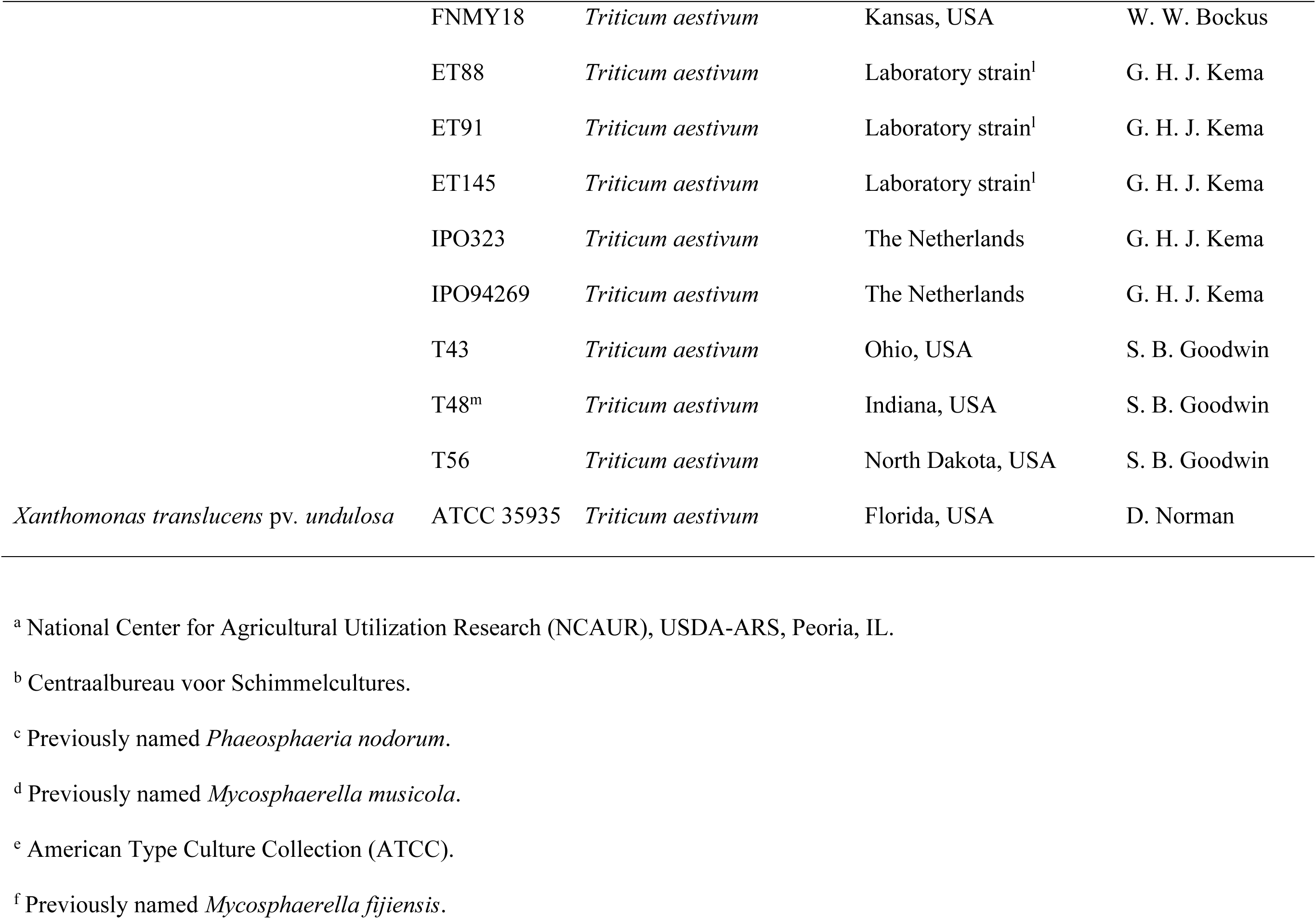
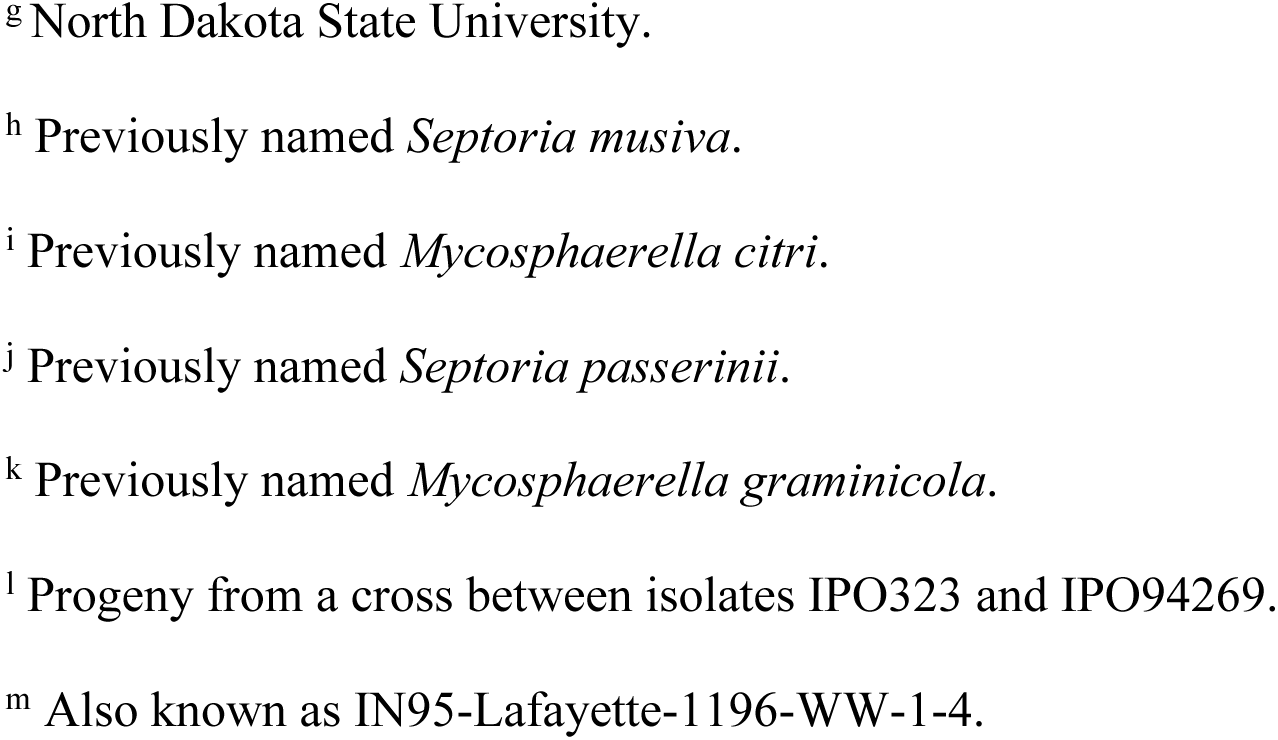
Fungal and bacterial species and isolates tested for the specificity of the real-time PCR primers developed for analysis of *Zymoseptoria tritici* DNA.

### Design and selection of species-specific primers

Two specific primer sets were designed between the intron and exon region of the ß-tubulin gene sequence using Primer Express software (version 1.5; Applied Biosystems, Foster City, CA). An additional primer set published previously (Fraaije et al. 2001) and two primer combinations from the internal transcribed spacer (ITS) region (White et al. 1990) were also tested (Table 2). To evaluate the primer sets for specificity, sensitivity, and product size, DNA of field isolates of *Z. tritici* collected from the Netherlands and the United States (California, Indiana, Kansas, Ohio, and North Dakota) and nine isolates of closely related species in the family Mycosphaerellaceae were examined (Table 1). Most of these fungi were chosen because they are close relatives of *Z. tritici* (Goodwin et al. 2001), so they are the most likely to give false positives in the amplifications. In addition, two unrelated fungi from rice (*Magnaporthe oryzae* and *M. salvinii*) and significant wheat pathogens (*Cochliobolus sativus,* ten species of the *Fusarium graminearum* complex, *Gaeumannomyces graminis* var. *tritici, Pyrenophora tritici-repentis, Parastagonospora nodorum, Puccinia graminis tritici, P. triticina, P. striiformis*) plus the bacterial leaf streak pathogen of wheat *Xanthomonas translucens* pv. *undulosa* were tested for primer specificity and sensitivity by conventional and real-time PCR. We further tested the hypothesis that the primer sites are not conserved in other fungal pathogens of wheat by *blastn* searches of GenBank and of complete genome sequences of the powdery mildew pathogen *Blumeria graminis, F. graminearum, P. graminis tritici, P. tritici-repentis, P. nodorum* and of all species in the rust order Pucciniales and the powdery mildew order Erysiphales.

**TABLE 2.**
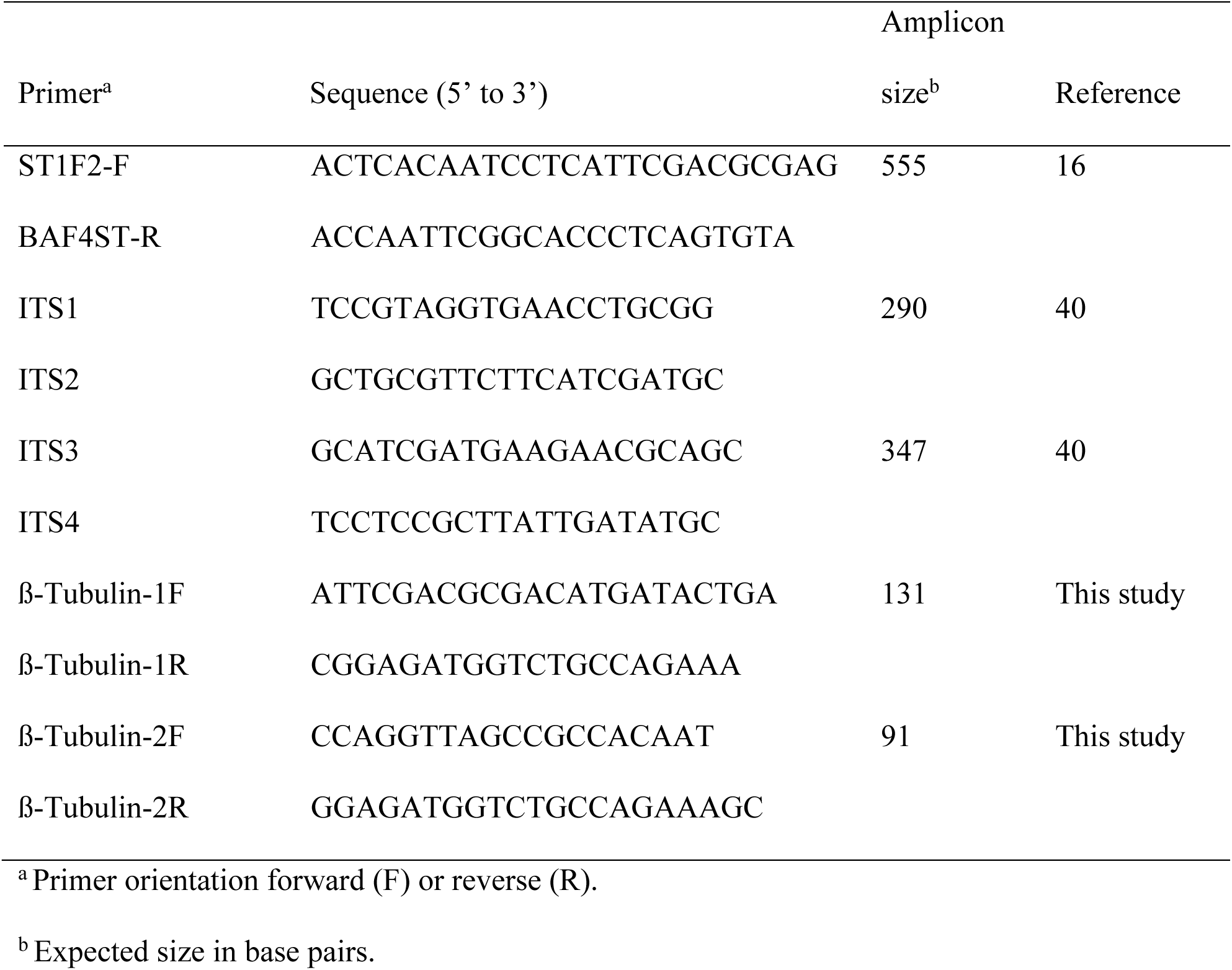
Sequences of primers used for the detection and quantification of *Zymoseptoria tritici* DNA with real-time quantitative PCR.

### Conventional and real-time PCR methods

For conventional PCR, each reaction contained 2 mM MgCl_2_, 200 μM of each dNTP, 0.2 μM of each primer, 12.5 ng of template DNA, and 1 unit of *Taq* DNA polymerase (Applied Biosystems, Foster City, CA) with distilled water added to make up 20 μl. Amplifications were performed in a PTC-100 Thermal cycler (MJ Research, Watertown, MA) at standard amplifications of 94°C for 2 min, followed by 35 cycles at 94°C for 30 s, 65°C for 1 min, and 72°C for 1.5 min, then with a final extension at 72°C for 4 min before cooling to 4°C. Products from each 20-µl reaction were separated by electrophoresis in 2% agarose (wt/vol) gels (Amresco, Solon, OH) at 4 V/cm in 0.5× TAE buffer (0.045 M Tris-acetate and 1 mM EDTA, pH 8.0). Validation experiments were performed before conducting actual experiments to select optimal concentrations of genomic DNA and primers for real-time PCR. These experiments analyzed a series of known amounts (1, 10, 20, 25, 50, and 100 ng) of genomic DNA isolated from inoculated plants of wheat cultivar Yecora Rojo. The concentrations of primers (0.25, 0.50, 1.25, 2.50, and 5.00 μM) also were optimized through a series of PCR tests. Optimal forward and reverse primer concentrations were selected, resulting in a minimum level of reporter fluorescence associated with an exponential increase in PCR product. Based on these results, 20 ng/μl of DNA and a 500-nM concentration of primers were selected for further experiments.

Real-time PCR was performed in MicroAmp® Optical 96-well reaction plates with barcodes using an ABI Prism 7700 Sequence Detection System (Applied Biosystems, Foster City, CA). Each real-time PCR reaction was in a total volume of 20 μl that contained 1× SYBR® Green PCR Master Mix (Applied Biosystems), 500 nM each of forward and reverse primers, and 20 ng of DNA template. The universal thermal-cycle protocol recommended by Applied Biosystems was used for PCR amplifications: pre-incubation at 50°C for 2 min, denaturation at 95°C for 10 min, followed by 45 cycles of denaturation at 95°C for 15 s, and annealing and extension together at 60°C for 1 min/cycle. Immediately after the final PCR cycle, a melting-curve analysis was done to determine the specificity of the primers by incubating the reaction at 95°C for 15 s, annealing at 60°C for 20 s, and then slowly increasing the temperature to 95°C over 20 min.

### Analyzing the relationship between disease severity and fungal DNA

To examine the relationship between disease severity and fungal DNA, pre-germinated seeds of two resistant (Tadinia and W7984) and two susceptible (Yecora Rojo and Opata 85) cultivars (Set 1) were transplanted into 10-cm-diameter plastic pots and grown in a greenhouse of the Department of Botany and Plant Pathology, Purdue University, West Lafayette, IN. Unless stated otherwise, each experiment was laid out in a randomized complete block design (RCBD) and repeated three times. The pots were arranged on a bench, and plants were inoculated with inoculum concentrations at 6 × 10^6^ spores/ml 40 days after sowing as described previously (Adhikari et al. 2003). Disease severities, expressed as the visually estimated percentage leaf areas of necrotic lesions (Adhikari et al. 2003; Gaunt et al. 1986; Rosielle 1972; Rosielle and Brown 1979), were scored at 1, 2, 3, 6, 9, 12, 15, 18, 21, 24, and 27 days after inoculation (DAI). Percent infected leaf area was used in this experiment because it was easier to see differences over time compared to pycnidia production. Three plants per pot and three pots per treatment at each time point were sampled using a destructive sampling method. After scoring for disease severity, the same plants were collected from each time point for DNA analysis. DNA from three plants from the same cultivar at each time point was pooled and analyzed by real-time PCR. Each DNA sample was analyzed three times by real-time PCR to estimate the assay variation.

### Quantifying the effects of low and high inoculum concentrations on fungal DNA

In a separate experiment, seeds of the two resistant (Tadinia and W7984) and two susceptible (Yecora Rojo and Opata 85) cultivars of Set 1 were planted as described above. Two inoculum concentrations, low (3 × 10^4^ spores/ml) and high (3 × 10^6^ spores/ml) were prepared, and plants were inoculated as described above. DNA from plants sampled at 6, 9, and 12 DAI was isolated and analyzed as independent biological replicates by real-time PCR. Remnant plants were maintained in the greenhouse for observation of symptoms. Each interaction (resistant or susceptible) was replicated over the two resistant and two susceptible cultivars.

### Estimating fungal DNA in segregating progenies

For this experiment, two resistant and two susceptible parents and the 10 resistant and 10 susceptible RILs of cultivar Set 2 were grown in pots and replicated three times. Plants were inoculated with ∼ 6 × 10^6^ spores/ml at the flag leaf stage. DNA from three plants of each RIL collected at 12 DAI was analyzed as independent biological replicates by real-time PCR. Estimating disease severity and fungal DNA on the same plants was not possible because they were sampled destructively before symptom expression. Therefore, disease scores, calculated as the average level of pycnidia production on a 0 to 5 scale within lesions on each leaf, were from previously published experiments (Adhikari et al. 2004a). Pycnidia production was used as the disease estimator in this experiment instead of the percent leaf area infected because it gave better discrimination between resistant and susceptible lines in previous tests (Kema et al. 1996). Remnant plants of each line were maintained in the greenhouse for an additional two weeks to test whether symptom development corresponded with the earlier classification.

### Quantifying fungal DNA at seedling and adult-plant stages in resistant and susceptible wheat cultivars

In this experiment, two resistant (Tadinia and Veranopolis) and two susceptible (Chinese Spring and Taichung 29) wheat cultivars (Set 3) were inoculated with ∼ 4 × 10^6^ spores/ml at 10 (for seedlings) and 40 (adult plants) days after sowing. At each time point, the two resistant and two susceptible cultivars were considered biological replications of resistance versus susceptibility. DNA was isolated from three plants from a single cultivar at 6, 9, and 12 DAI for each treatment and analyzed as independent biological replicates by real-time PCR. Remnant plants of each line were maintained in the greenhouse for another two weeks to verify symptom expression.

### Statistical analyses

A standard curve was constructed using known quantities (0.006, 0.024, 0.098. 0.39, 1.56, 6.25, 25, and 100 ng) of DNA from *Z. tritici* (isolate T48) amplified with the primer set ß-tubulin-2. For each reaction, C_T_ values were plotted against the log_10_ of the initial starting quantity of DNA. By comparison of the C_T_ values against the log DNA amount of *Z. tritici*, a linear regression was calculated and used to quantify the DNA. In all experiments, this standard curve was used to calculate fungal DNA in infected wheat cultivars based on the C_T_ value of the test sample, as revealed by real-time PCR. For each unknown sample, 20 ng of DNA template from inoculated wheat leaves were loaded in a real-time PCR reaction, and fungal DNA was quantified in nanograms (ng) based on the C_T_ value of the sample. To test whether plant DNA affected fungal DNA estimation, additional standard curves were made with pure fungal DNA (0.001, 0.005, 0.020, 0.078, 0.313, 1.25, 5, and 20 ng) alone and spiked with plant DNA to make up 20 ng in each reaction (e.g., for the spiked test, the sample with 1.25 ng of fungal DNA contained 20 - 1.25 = 18.75 ng of plant DNA).

DNA yield and disease severity data obtained from each cultivar or line at each time point were subjected to an analysis of variance (ANOVA) using the general linear model procedure of the Statistical Analysis System (SAS Institute, Cary, NC). Arcsin-square root transformations of the disease severity data were made before analysis. Means were separated and differences were estimated using linear contrasts. Spearman rank correlations were used to determine the relationships between disease severity and fungal DNA after a log_e_ transformation of the data to improve the homogeneity of variances. Data from the two resistant and two susceptible cultivars were analyzed individually or combined to represent the biological replication of each interaction (resistant or susceptible) at the same point in time. Standard deviations of the means were calculated at specific time points. A pair-wise *t-*test was used to determine if differences in disease severity or fungal DNA were significant (*P* ≤ 0.05) between the means of the resistant and susceptible cultivars.

## Results

### Primer specificity and detection of *Z. tritici* in wheat cultivars by conventional PCR

To evaluate the specificity of the primer pairs, the ITS primer combinations generated products from both the host and the pathogen (Fig. 1A), confirming that these primers did not have the high specificity required for the real-time PCR method. The primer pairs ß-tubulin-1 and ß-tubulin-2 produced a single amplicon of approximately 130 or 90 bp (Fig. 1A) from the genomic DNA of several field isolates and progeny of *Z. tritici*. Primer pairs ß-tubulin-1 (*data not shown*) and ß-tubulin-2 (Fig. 1B) did not amplify DNA from the non-target barley pathogen *Z. passerinii*. Because it gave a smaller product, the primer pair ß-tubulin-2 was chosen for all subsequent analyses. No amplification was detected with primer pair ß-tubulin-2 from uninoculated (healthy) wheat leaves (Fig. 1A), but DNA from all isolates of *Z. tritici* tested was amplified (Fig. 1B). The specificity of this primer set was confirmed by the absence of conventional PCR products when tested on genomic DNA from a wide range of fungal pathogens of wheat plus related species in the same fungal family (Table 1). Amplification was only observed with *Z. tritici* isolates and the very closely related *S. triseti* isolated from *Phalaris arundinacea*, which should be reclassified into the genus *Zymoseptoria* (Fig. 1C) (S. B. Goodwin, *unpublished data*). No amplification was detected in PCR amplifications of DNA from the other fungal and bacterial pathogens of wheat and other hosts tested (Figs. 1C and 1D). The *blastn* searches of the GenBank non-redundant database and on the complete genome sequences for *F. graminearum, P. graminis tritici, Pyrenophora tritici-repentis*, and *Parastagonospora nodorum* (http://www.broadinstitute.org/science/projects/fungal-genome-initiative and http://www.jgi.doe.gov/genome-projects) plus all fungi in the orders Pucciniales and Erysiphales further confirmed that the primer sites were absent in these fungi. The ß-tubulin-2 forward primer had *blastn* matches to ß-tubulin sequences from isolates of *Z. tritici* but not to those from any other fungi. The sequence of the ß-tubulin-2 reverse primer was more conserved, with identical matches to several fungi, mostly various species of *Zymoseptoria* but also including two species of *Fusarium*. Sequences of *F. graminearum* and *P. nodorum* matched the ß-tubulin-2 reverse primer at 20 of the 21 bases (*data not shown*). However, the complete lack of similarity of the ß-tubulin-2 forward primer would prevent amplification with DNA from these species even if the reverse primers did not have a single-base mismatch.

**Fig. 1.**
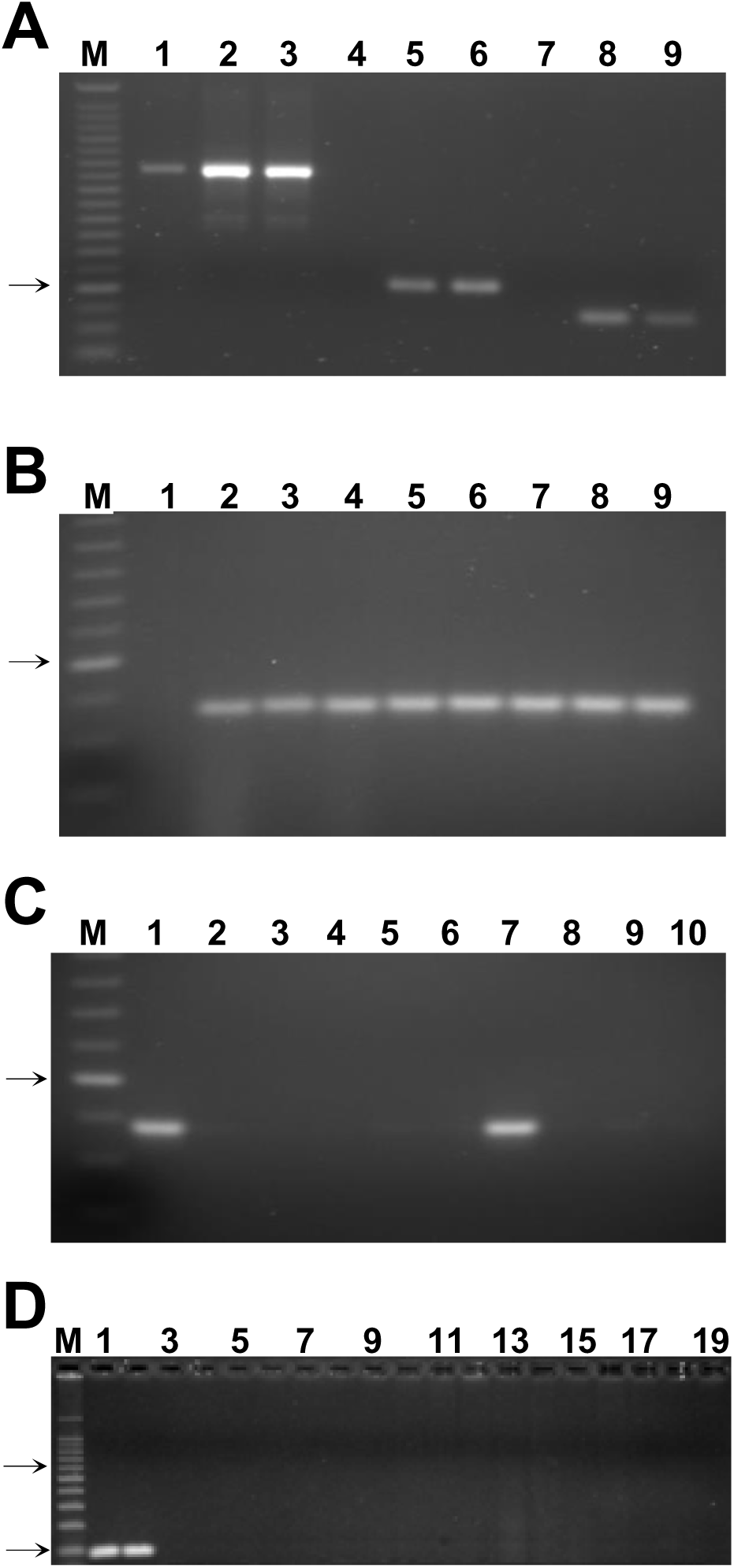
Genomic DNA analyzed by conventional polymerase chain reaction (PCR). **A,** Genomic DNA was isolated from the susceptible wheat cultivar Yecora Rojo uninoculated at 1 and 24 days after inoculation with Indiana isolate T48 of *Zymoseptoria tritici* in the greenhouse. PCR products were amplified with: lanes 1-3, the internal transcribed spacer (ITS) primers ITS1 and ITS2; lanes 4-6, primer pair ß-tubulin-1; lanes 7-9, primer pair ß-tubulin-2; and separated in a 2% (wt/vol) agarose gel. Lanes: M, 25-base pair (bp) ladder; 1, 4, and 7, Yecora Rojo uninoculated (control); 2, 5, and 8, Yecora Rojo 1 day after inoculation; 3, 6, and 9, Yecora Rojo 24 days after inoculation. **B**, PCR products of field isolates of *Z. tritici* and progeny DNA amplified with the ß-tubulin-2 primer set. Lanes: M, 25-bp ladder; 1, *Zymoseptoria passerinii* isolate P64; 2, *Z. tritici* isolate T56; 3, *Z. tritici* isolate T43; 4, *Z. tritici* isolate T46; 5, *Z. tritici* isolate IPO323; 6, *Z. tritici* isolate 94269; 7, *Z. tritici* isolate ET88; 8, *Z. tritici* isolate ET91, and 9, *Z. tritici* isolate ET145. **C**, PCR products of DNA from different fungal species amplified with the ß-tubulin-2 primer set. Lanes: M, 25-bp ladder; 1, *Z. tritici* isolate T48; 2, *Mycosphaerella punctiformis*; 3, *M. macrospora*; 4, *Pseudocercospora musae*; 5, *Zasmidium citri-grisea*; 6, *P. fijiensis*; 7, *Septoria triseti*; 8, *S. lycopersici*; 9, *Shaerulina musiva*; and 10, *Magnaporthe salvinii*. The arrow on the left of figures 1A-C indicates the 125-bp band of the 25-bp size-standard ladder. **D**, PCR products from DNA of fungal and bacterial isolates tested for specificity with the ß-tubulin-2 primer set. Lanes: M, 100-bp ladder; 1, *Z. tritici* isolate CA16 (from California); 2, *Z. tritici* isolate FNMY 18 (from Kansas); 3, *Puccinia triticina* isolate 2005-L; 4, *P. striiformis* isolate 2k-41-Yr9 (race PST-78); 5, *P. graminis* f.sp. *tritici* isolate 2005-S2; 6, *Cochliobolus sativus* isolate CS45; 7, *Parastagonospora nodorum* isolate Sn2000; 8, *Pyrenophora tritici-repentis* isolate Pti2 (race 1); 9, *Fusarium meridonale* isolate NRRL 28436; 10, *F. acacia-mearnsii* isolate NRRL 26754; 11, *F. mesoamericanum* isolate NRRL 25797; 12, *F. asiaticum* isolate NRRL 13818; 13, *F. austroamericanum* isolate NRRL 2903; 14, *F. culmorum* isolate NRRL 25475; 15, *F. pseudograminearum* isolate NRRL 28334; 16, *F. cerealis* isolate NRRL 1372; 17, *F. boothi* isolate NRRL 26916; 18, *F. graminearum* isolate NRRL 31084; and 19, *Xanthomonas translucens* pv. *undulosa* strain ATCC 35935. Only odd-numbered lanes are labeled. The top and bottom arrows to the left of Fig. 1D indicate the positions of the 500- and 100-bp size standards, respectively, in marker lane M.

### Sensitivity and host-versus-pathogen specificity of the real-time PCR method

Melting-curve analysis provided a graphical representation of the PCR product after amplification. A peak within the melting-temperature range of 80 to 85°C (Fig. 2A) revealed that the ß-tubulin-2 primer pair amplified a specific PCR product without interference from primer dimers or host DNA. Amplification profiles were similar over the range of DNA template quantities from 0.006 to 100 ng (Fig. 2B). A standard curve using genomic DNA of the isolate of *Z. tritici* used for greenhouse inoculations was constructed (Fig. 2C). The correlation coefficient (*r* = -0.997) between the log_10_ of the initial DNA quantity and the C_T_ value was highly significant (*P* < 0.0001), suggesting that the individual data points were highly correlated with the calculated linear regression. As expected, no amplification was detected with DNA from healthy (uninoculated) wheat leaves (*data not shown*). Spiking the samples with plant DNA flattened the standard curve slightly at the lowest pathogen DNA concentrations and increased the variance of some of the measurements (*data not shown*). However, because the standard curves were virtually identical within the observed range of pathogen DNA quantities (tested by paired t-tests), a pathogen standard curve was used for all estimates of fungal DNA.

**Fig. 2.**
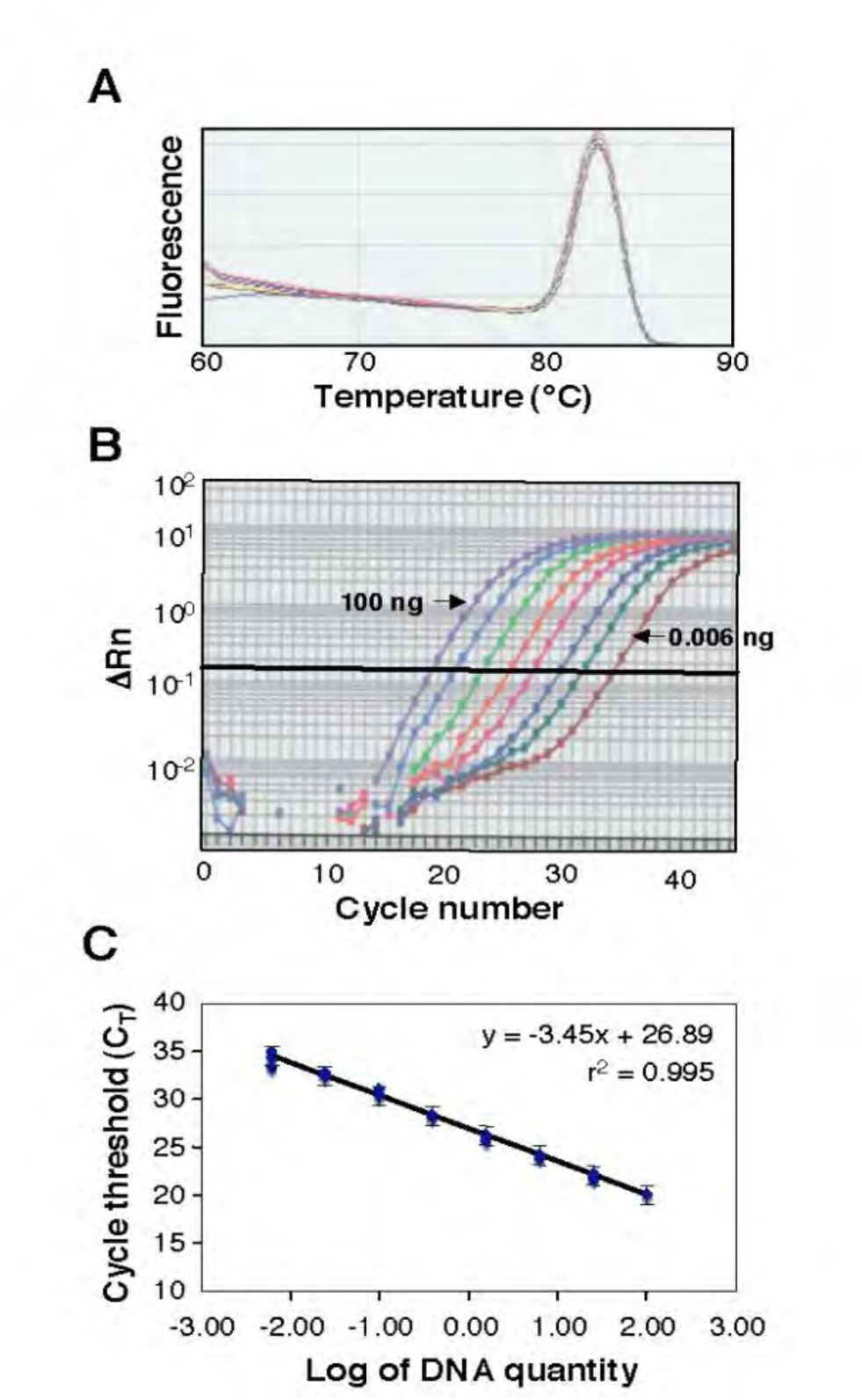
A melting curve constructed from a real-time polymerase chain reaction (PCR) analysis. **A**, Genomic DNA was isolated from the susceptible wheat cultivar Yecora Rojo 24 days after inoculation with *Zymoseptoria tritici* isolate T48. For each reaction, 20 ng of DNA was amplified with the primer set ß-tubulin-2. **B**, Real-time amplification profiles of *Z. tritici* DNA samples generated from cycle-by-cycle collection of fluorescent light emission (ΔRn). **C**, A standard curve using eight known DNA quantities (0.006, 0.024, 0.098, 0.39, 1.56, 6.25, 25, and 100 ng) from *Z. tritici* amplified with the primer set ß-tubulin-2. Three replicate PCR reactions were performed for each quantity. A regression equation was calculated from the standard curve and used for the conversion of C_T_ values to the fungal DNA amount. Brackets indicate the standard errors.

### Assessing the relationships between disease severity and fungal DNA

Necrotic lesions were visible, and pycnidia developed beginning at 16 or 18 DAI on the susceptible cultivars Opata 85 and Yecora Rojo, respectively. Disease severity progressed rapidly afterward and reached about 40 to 70% in both susceptible cultivars by 27 DAI (Fig. 3A). In contrast, no significant necrosis or discoloration was observed in the two resistant cultivars Tadinia and W7984, even at 21 DAI. There was a slight increase of disease severity in W7984 between 21 and 27 DAI (Fig. 3B). The susceptible cultivars had significantly more disease than the resistant cultivars at days 21 and 27, and the difference was nearly significant (*P* = 0.0524) at day 24.

**Fig. 3.**
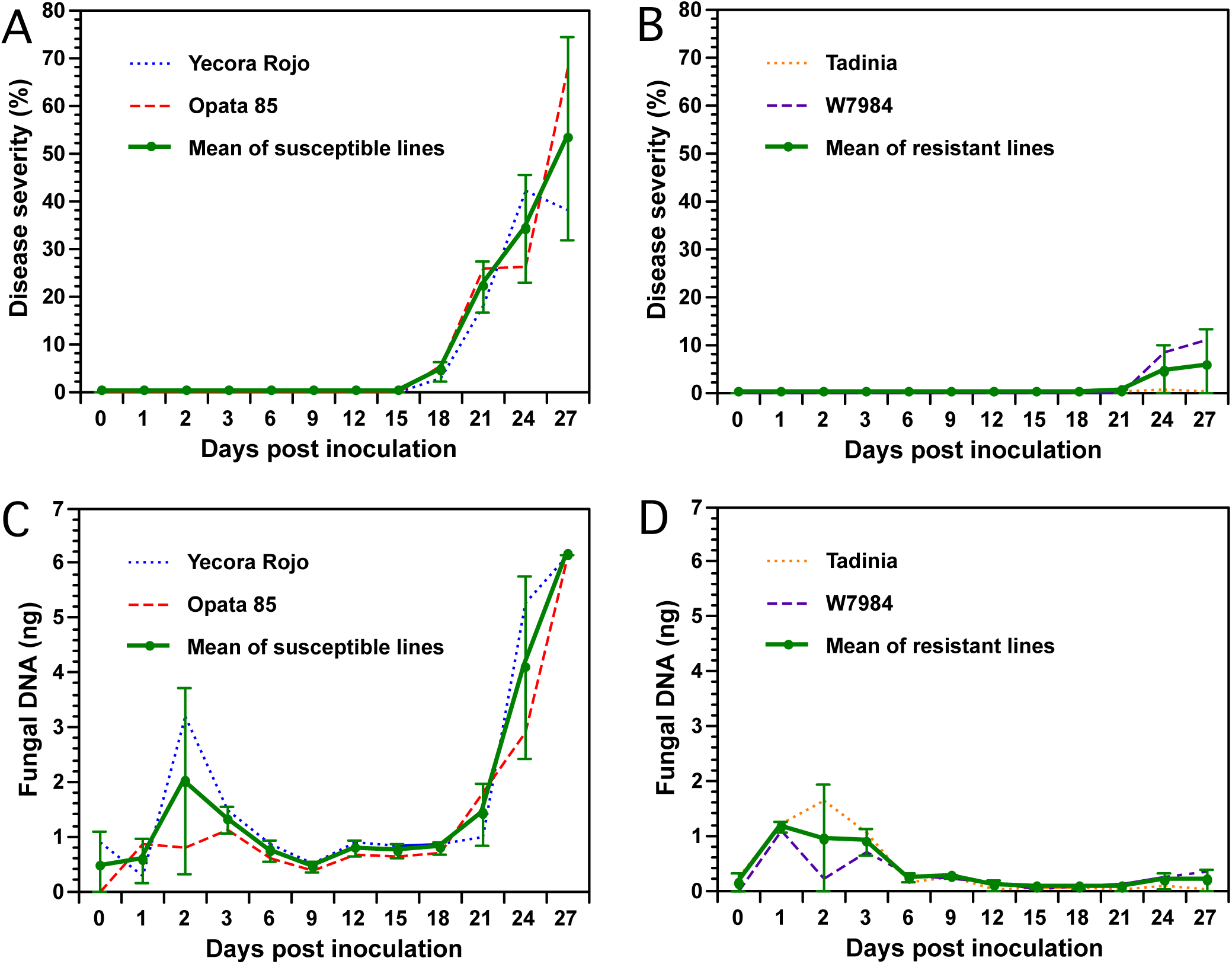
Determination of Septoria tritici blotch (STB) progression and fungal DNA in two resistant (Tadinia and W7984, containing the *Stb4* and *Stb8* genes, respectively) and two susceptible (Opata 85 and Yecora Rojo) wheat cultivars inoculated with *Zymoseptoria tritici* isolate T48 in a greenhouse. **A**, Progression of STB in the two susceptible cultivars. **B,** Progression of STB in the two resistant cultivars. Disease severities (DS) in Figs. 3A and 3B were expressed as percentage leaf area of necrotic lesions for each cultivar. Each data point represents the mean of three replicate plants. **C**, DNA of *Z. tritici* in the two susceptible cultivars as measured by real-time polymerase chain reaction (PCR). **D**, DNA of *Z. tritici* in the two resistant cultivars. Each sample for real-time PCR in Figs. 3C and 3D contained pooled leaves collected from three inoculated plants grown in individual plastic pots. Each data point represents the average of three PCR analyses (technical replications) of the same DNA sample to estimate the assay variance. Data for each cultivar and the mean over both cultivars are plotted separately on each figure. Error bars indicate the 95% confidence intervals of the means of the two susceptible or two resistant cultivars averaged at each time point.

Fungal DNA estimated by real-time PCR increased slightly in the resistant and susceptible cultivars during the first 2 or 3 DAI (Figs. 3C and 3D). However, fungal DNA remained low from 6 to 9 DAI and was not significantly different between resistant and susceptible wheat cultivars. Data analysis revealed that significantly more fungal DNA was detected in the two susceptible cultivars beginning 12 DAI compared to the two resistant cultivars (Figs. 3C and 3D). Fungal DNA remained constant in all cultivars between 12 and 18 DAI but increased almost exponentially in both susceptible cultivars following the development of necrosis and pycnidial production beginning 18 or 21 DAI. In contrast, no fungal or a low level of fungal DNA was detected in the resistant cultivars Tadinia and W7984 between 12 and 21 DAI. Interestingly, fungal DNA did increase slightly in resistant cultivar W7984 between 21 and 27 DAI (Fig. 3D), mirroring the slight increase in disease severity (Fig. 3B). Pair-wise comparisons of means using *t-*tests indicated that the differences for fungal DNA between the resistant and susceptible wheat cultivars were significant at *P <* 0.0001 from 12 to 18 and at 27 DAI, and at *P* < 0.08 from 21 to 24 DAI. The Spearman rank correlations between the mean disease severities and the amounts of fungal DNA detected in the susceptible cultivars Yecora Rojo (R^2^ = 0.93) and Opata 85 (R^2^ = 0.92) were positive and significant (*P <* 0.0001).

### Determining the effects of low and high inoculum concentrations on fungal DNA

No differences in fungal DNA were detected with the low inoculum concentration. Although the estimated fungal DNA was slightly higher in the susceptible cultivars at 12 DAI the difference was not significant (Fig. 4A). In contrast, large and significant differences in fungal DNA (*P <* 0.0001) were detected at 12 DAI with the high inoculum concentration between the resistant cultivars Tadinia and W7984 compared to the susceptible cultivars Yecora Rojo and Opata 85 (Fig. 4B). No significant differences in fungal DNA were observed either between the two resistant cultivars or the two susceptible cultivars. The remnant plants of the susceptible cultivars all showed symptoms of STB, but disease in those inoculated with the lower spore concentration was delayed by several days (*data not shown*).

**Fig. 4.**
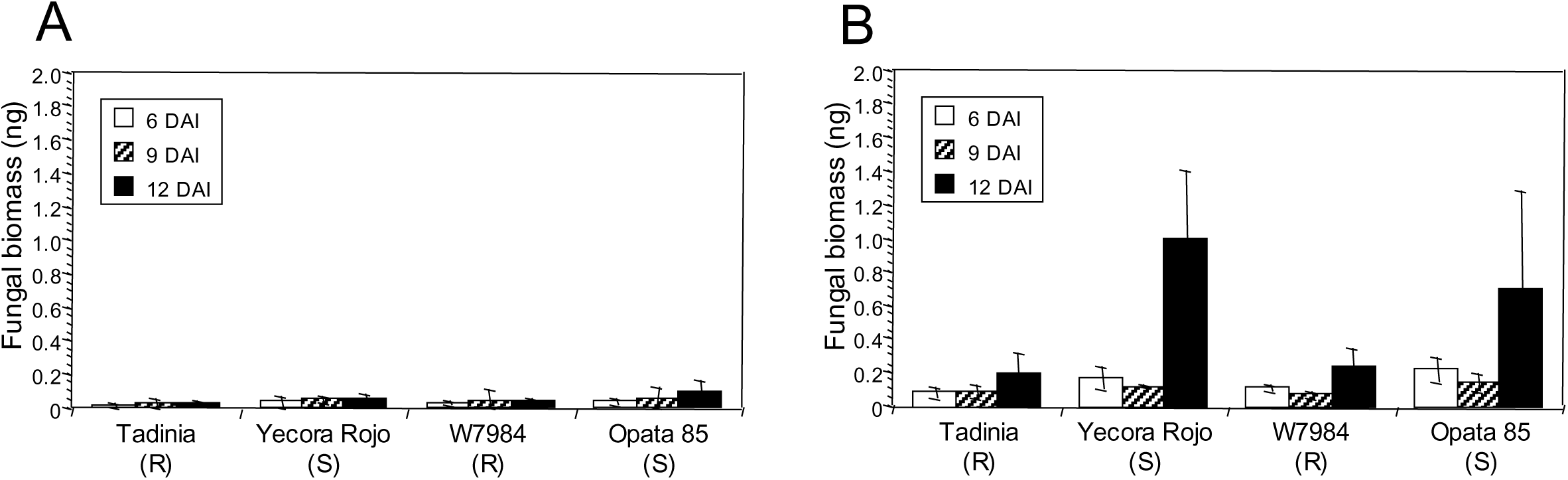
Effect of inoculum concentration on estimation of DNA of *Zymoseptoria tritici* in resistant and susceptible wheat cultivars. Two resistant (Tadinia and W7984) and two susceptible (Opata 85 and Yecora Rojo) cultivars of wheat were inoculated 13 days after sowing with (**A**) 3 × 10^4^ or (**B**) 3 × 10^6^ spores/ml of *Z. tritici* isolate T48. Three plants of each treatment were collected separately as biological replicates 6, 9, and 12 days after inoculation (DAI), and fungal DNA (ng) was estimated by real-time PCR. Standard deviations of the means are indicated by error bars above each histogram entry.

### Estimating fungal DNA in progenies segregating for resistance

Six resistant RILs (1703, 1730, 1743, 1753, 1767, and 1779) showed no symptoms during previous testing (Adhikari et al. 2004a). In contrast, extremely low pycnidial densities (0.33 to 0.5) had been observed on the other four resistant RILs (1707, 1712, 1740, and 1760) (Fig. 5). All susceptible RILs had high pycnidial densities (3.67 to 4.83) in previous testing (Adhikari et al. 2004a). Results obtained with real-time PCR showed a strong relationship between estimates of *Z. tritici* infection and fungal DNA. Less fungal DNA was detected in the resistant cultivar Tadinia and the 10 resistant RILs (0.21 to 0.71 ng) than in the susceptible cultivar Yecora Rojo or the susceptible RILs (0.80 to 5.23 ng) (Fig. 5), with only four exceptions. The amounts of fungal DNA for Yecora Rojo and four susceptible RILs (1722, 1725, 1727, and 1759) were not significantly higher than those for the resistant RILS with the highest levels of fungal DNA, although they were higher than the mean of all resistant RILs. All resistant RILs had DNA estimates of 0.71 ng or lower, while all susceptible RILs had values of 0.80 or higher, so a value of 0.75 ng discriminated between resistance and susceptibility. It was not possible to estimate disease severity and fungal DNA in the same plants due to destructive sampling for DNA extraction before symptom expression at 12 DAI, but the DNA results gave good agreement with previous phenotypic testing.

**Fig. 5.**
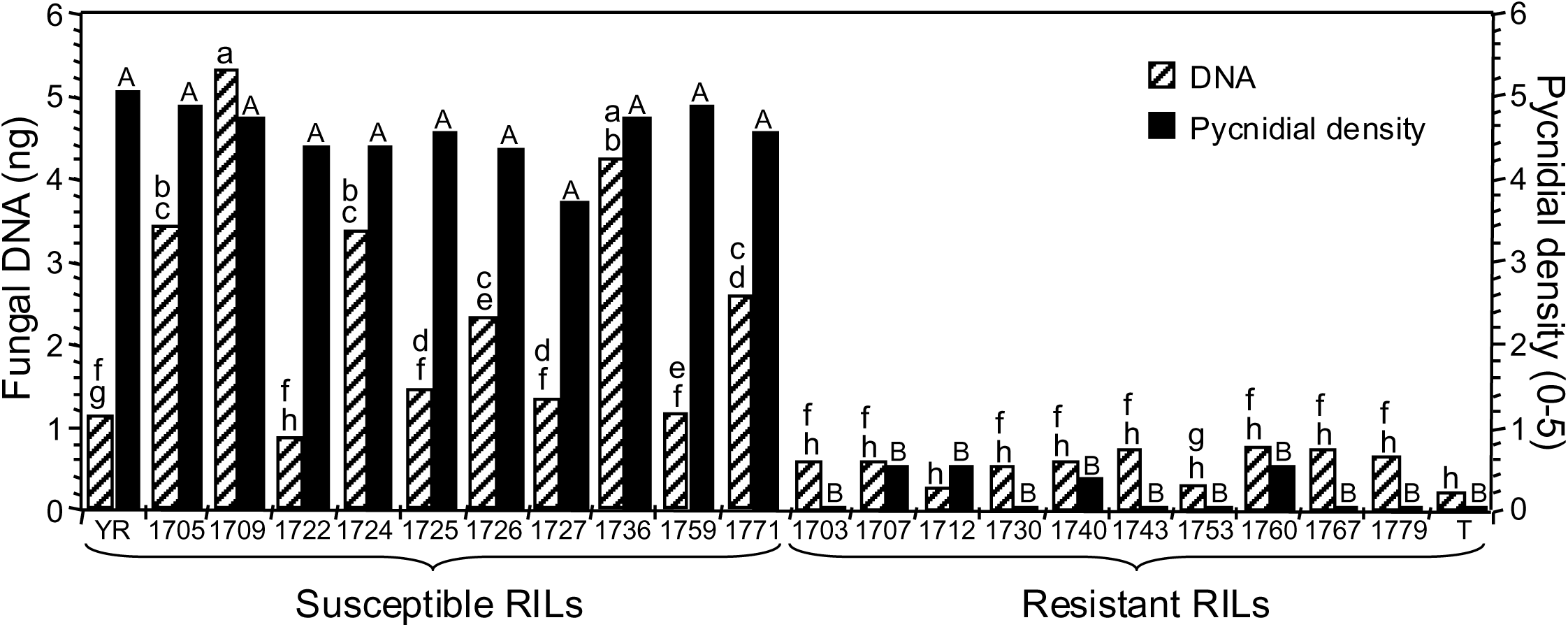
The relationship between *Zymoseptoria tritici* DNA at 12 days after inoculation (DAI) estimated with real-time PCR and phenotypic disease scores at 28 DAI based on pycnidial density within lesions in 10 resistant and 10 susceptible recombinant-inbred lines (RILs) from a resistant × susceptible wheat cross. Generation of the RILs, phenotypic disease scoring, and mapping of the *Stb4* gene for resistance were published previously (3). YR, susceptible parent Yecora Rojo; T, resistant parent Tadinia (*Stb4* gene); resistant and susceptible RILs are indicated by number. Histogram bars represent the mean of six plants treated as biological replicates; those with different letters are significantly different for fungal DNA (lower case) or pycnidial density (small capitals) according to Fisher’s protected least significant difference (*P* < 0.05). A range is indicated by providing the first and last letters.

### Quantifying fungal DNA at seedling and adult-plant stages in resistant and susceptible wheat cultivars

At both growth stages, significantly less fungal DNA (*P <* 0.0001) was detected in the resistant cultivars Tadinia and Veranopolis than in the susceptible cultivars Chinese Spring and Taichung 29 (Fig. 6). Chinese Spring had significantly more fungal DNA compared to Taichung 29 at 12 DAI in the seedling test (Fig. 6A), but no significant differences in the amounts of pathogen DNA were observed either between the two resistant cultivars or the two susceptible cultivars in adult plants (Fig. 6B). The estimated levels of fungal DNA were higher in the susceptible cultivars in adult compared to seedling tests.

**Fig. 6.**
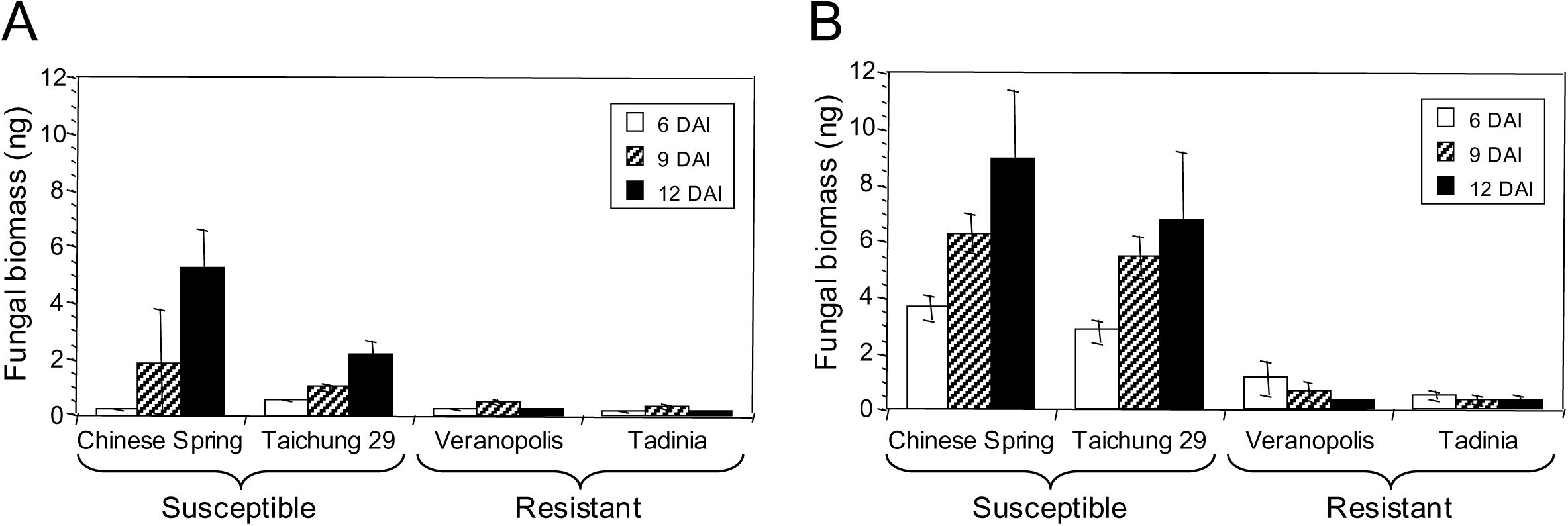
DNA of *Zymoseptoria tritici* in the two resistant cultivars Tadinia (*Stb4* gene for resistance) and Veranopolis (*Stb2*) and the two susceptible cultivars Chinese Spring and Taichung 29 at (**A**) seedling and (**B**) adult-plant stages as measured by real-time PCR. Fungal DNA was determined at 6, 9, and 12 days after inoculation with isolate T48 in the greenhouse. Each data point represents the average and standard deviation of three independent biological replications from a single cultivar at a specific time.

## Discussion

In these experiments, specific primers were designed for a particular region of the ß-tubulin gene that provided an efficient method to observe the changes in the amount of *Z. tritici* DNA during compatible and incompatible interactions in wheat leaves. Previous primers for multiplex and real-time quantitative PCR of *Z. tritici* based on sequences of the ß-tubulin gene (Fraaije et al. 2001; Guo et al. 2006) were specific for *Z. tritici* when tested on a range of unrelated fungi that also cause disease in wheat. Still, they gave a product of 555 bp, outside the 50-150-bp range recommended for real-time PCR. In addition, those primers were not specific for *Z. tritici* when tested against related *Septoria, Mycosphaerella* and *Zymoseptoria* species. A more elegant approach used the real-time PCR method and TaqMan probes to detect specific alleles of the cytochrome *b* gene for resistance or susceptibility to fungicides (Fraaije et al. 2005). Although that method can monitor fungal DNA (Keon et al. 2007), it is more complicated and expensive than the SYBR green technique. This newly developed real-time PCR method can accurately detect and quantify *Z. tritici* DNA to estimate disease resistance in wheat cultivars.

The primer set ß-tubulin-2 was highly specific for *Z. tritici* isolates except for *S. triseti*, the most closely related to *Z. tritici* of the species tested (S. B. Goodwin, *unpublished data*), and which therefore should be reclassified into the genus *Zymoseptoria*. The isolate of *S. triseti* tested was from *Phalaris arundinacea* (reed canarygrass), and this pathogen is specific for that host, so cross amplification should not be a problem when the ß-tubulin-2 primers are used on infected wheat. More importantly, no amplification was detected from the genomic DNA of key fungal and bacterial pathogens of wheat and the other *Septoria* and *Mycosphaerella* species tested from other hosts. Other major fungal pathogens of wheat are too distantly related for the primer sites to be conserved, as confirmed by *blastn* analyses of GenBank and genomic sequences, including all members of the Pucciniales and Erysiphales. These results suggest that the primer set ß-tubulin-2 was highly specific for use in the real-time PCR assay. High negative correlations between the DNA quantities estimated from standard curves and the C_T_ values further suggest that this method is reliable. In addition, no amplification was detected in DNA isolated from healthy or un-inoculated wheat leaves, confirming that the primer set ß-tubulin-2 was specific for *Z. tritici*.

There were notable changes in the amounts of fungal DNA found in the susceptible cultivars throughout the 27-day time course. Initially, during the first 3 DAI in both resistant and susceptible cultivars, there was an increase in fungal DNA, which was likely due to spore germination and epiphytic growth of the fungus in the enclosed inoculation chamber. This early stage of pathogenesis is essential for establishing disease, spreading the fungus to the apoplast, and colonizing neighboring sub-stomatal cavities (Duncan and Howard 2000; Fones et al. 2017). Subsequently, there was a gradual decrease in fungal DNA from about 2 or 3 DAI to 6 or 9 DAI. This decrease was probably due to a change in conditions from the sheltered environment of the inoculation chamber to the relatively high solar radiation and low humidity of an unprotected greenhouse bench at 3 DAI. Any fungal hyphae remaining outside the host at this stage were likely to die rapidly, which helps explain the observed decrease in fungal DNA.

During the latent phase of infection, small but statistically significant differences in fungal DNA were detected between the resistant and susceptible cultivars beginning at 12 DAI. The level of fungal DNA remained relatively consistent and low in the susceptible cultivars until symptoms became evident at around 18 DAI. After that, both symptoms and fungal DNA increased dramatically. Therefore, there was no significant increase in fungal DNA during the latent period of the disease. A similar pattern occurred with the rust pathogen *P. graminis* f. sp. *tritici*, in which the increase in fungal DNA in susceptible interactions was concomitant with symptom expression (Mayama et al. 1975). However, this result contrasts with those from fungi such as the rice blast pathogen, *M. oryzae*, which showed a rapid and constant increase in fungal DNA beginning two days after inoculation (Qi and Yang 2002). The data for *Z. tritici* suggest that a developmental switch may be initiated to cause the rapid increase in fungal DNA and onset of symptoms. This switch could involve toxins such as those produced by some species related to *Z. tritici* (Perrone et al. 2001). For example, cercosporin is created by members of the anamorphic genus *Cercospora* (Perrone et al. 2001). Another hypothesis is that the pathogen triggers programmed cell death in the host (Keon et al. 2007). Whatever the mechanism, these dynamic changes in *Z. tritici* DNA in the susceptible cultivars were significantly correlated with the onset of symptoms so can serve as an indicator of susceptibility. Similarly high correlations between disease susceptibility and *Z. tritici* DNA estimated by real-time PCR were reported by Guo et al. (2006), but that analysis did not include resistant cultivars.

Intriguingly, it was observed that there was a slight increase in disease severity and fungal DNA in the resistant cultivar W7984 at 24 DAI. These results support the previous findings that *Z. tritici* can cause infection in incompatible interactions, but at a much lower level than in compatible interactions (Chu et al. 2019; Kumar et al. 2015; Milgate et al. 2023; Oliver et al. 2008; Ozdemir et al. 2020; Ware et al. 2003). Even though such a low level of fungal infection may not be significant for disease epidemiology, it could still help maintain genetic diversity within populations. For instance, if incompatible strains can fertilize compatible members of the opposite mating type, it could initiate the sexual cycle.

A PCR-based method for quantifying *Z. tritici* DNA in plants is highly desirable due to the long latent period of STB disease. Previous studies have used disease severity and pycnidial density to identify cultivars with increased resistance to *Z. tritici* (Eyal et al. 1987; Goodwin 2007; Brown et al. 2015; Ors et al. 2018). Evaluating genetic materials for resistance to *Z. tritici* under greenhouse and field conditions requires visual assessment, which usually takes 24 to 28 days after inoculation. In addition to the genetic component, the environment during the time of symptom expression, variation among evaluators, and use of an integer scale for disease rating (Gaunt et al. 1986; Rosielle and Brown 1979) also can affect the accuracy of disease scoring and complicate the analysis and interpretation of data. On the other hand, using a specific ß-tubulin-2 primer set in real-time PCR at 12 DAI provides a less subjective estimate of disease severity, making it a more reliable method. However, differences in fungal DNA between resistant and susceptible cultivars at earlier time points were minor. Therefore, the real-time PCR approach, in its current form, is best suited for use in a secondary role in a breeding program to minimize the effect of environmental variations on disease expression and to provide an unbiased estimator of quantitative resistance. This study analyzed two resistant wheat cultivars, Tadinia and W7984, which possess specific resistance genes *Stb4* and *Stb8*, respectively. These genes have been mapped to chromosomes 7D (Adhikari et al. 2004a) and 7BL (Adhikari et al. 2003) and probably were contributed by wheat’s B- and D-genome ancestors. Despite their separate evolutionary origins, both had a similar effect on fungal DNA accumulation during pathogenesis. This likely extends to most other STB resistance genes. Similar strong correlations between visual estimates of disease severity and DNA quantity also were shown for *Z. tritici* in bread (Gouache et al. 2009) and durum (Tonti et al. 2019) wheat.

Real-time PCR was used to distinguish between wheat cultivars and correctly segregated RILs into resistant and susceptible classes based on disease symptoms. The cultivars and RILs rated as susceptible to *Z. tritici* had more fungal DNA in their leaves than the resistant ones. In contrast, the lowest fungal DNA was detected in the resistant cultivars and RILs. The Spearman rank correlations between pycnidial density and fungal DNA were positive and significant for both susceptible cultivars. However, four RILs and Yecora Rojo had higher fungal DNA than the resistant RILs, but the difference was insignificant. It is unclear why this discrepancy occurred, but it could be due to biological or operational factors. One possible explanation is that the plants with lower-than-expected fungal DNA received less inoculum, possibly because they grew more slowly than the other lines tested.

The pycnidial density scores were calculated based on several years of testing of adult plants, which were scored at 24 to 28 DAI (Adhikari et al. 2003; Adhikari et al. 2004a). Plant samples for real-time PCR analysis were collected at 12 DAI, the earliest point at which significant differences were observed before the exponential increase in biomass seen in the time-course experiment. Since the plants were collected for DNA analysis before symptom expression, it was impossible to know whether they would have shown the same pycnidial densities as the same lines when tested as adult plants scored at 28 DAI. This uncertainty could result in some susceptible RILs being scored as resistant based on real-time PCR results alone. Delaying the sample collection by one or two days could eliminate this problem. However, the opposite error of resistant plants falsely scored as susceptible did not occur in our tests. Our results were consistent with those obtained with other previously described methods to quantify fungal growth in wheat cultivars (Chu et al. 2019; Cohen and Eyal 1993; Keon et al. 2007; Oliver et al. 2008; Pnini-Cohen 2000; Stergiopoulos et al. 2003). Highly positive and significant correlations between fungal DNA and disease severity also were observed in alfalfa to *Aphanomyces euteiches* (Vandermark et al. 2002), in wheat to *Fusarium* spp. (Kumar et al. 2015; Milgate et al. 2023; Ozdemir et al. 2020), and with several pathogens of *Arabidopsis* (Brouwer et al. 200; Gachon and Saindrenan 2004).

Our analyses revealed that there were more noticeable differences in fungal DNA when the tests were conducted on adult plants instead of at seedling stages. However, even the seedling tests provided good discrimination. One explanation for this result may be that adult plants provide a better substrate for fungal growth, and thus, more significant differences in fungal DNA yield. Another reason may be how the plants were harvested. In the seedling tests, the entire small plant was clipped off at the soil level, so the samples included the stems that typically do not show symptoms, plus leaf tissue that grew after the plant was inoculated. These may have decreased the amounts of fungal DNA relative to the total samples. In contrast, only the leaves were harvested from the much larger adult plants, and no additional leaves grew after inoculation, so these samples included only tissue that was inoculated. Excluding stems and the newest leaves from the seedling samples might better represent the actual level of fungal DNA and provide a more accurate separation between resistant and susceptible plants. Inoculum concentration also was critical, as no differences were apparent at the lower inoculum concentration by 12 DAI. Symptoms did appear on these plants, but they were delayed by several days relative to those inoculated with more spores. Therefore, delaying the assay by two or three days or increasing the concentration of the inoculum beyond the highest level tested in our experiments may lead to increased discrimination between resistant and susceptible wheat lines.

The real-time PCR assay also should be tested with different pathogen isolates. Variation in virulence (Eyal et al. 1985; Kema et al. 1996) and specific adaptation by *Z. tritici* to vertically resistant wheat cultivars (Cowger et al. 2000; Jackson et al. 2000; Ababa and Mekonnen 2024) may affect the fungal DNA produced by particular cultivar-isolate combinations. Accordingly, it is necessary to investigate the amount of fungal DNA in planta that is proportional to the development of disease symptoms induced by different virulent isolates or pathotypes of *Z. tritici*. A potential approach to enhance the technique could be to use a plant gene as a reference for normalization and endogenous control (Valseia et al. 2005; Winton et al. 2002). In this case, fungal DNA would be estimated as a percentage of the total amplifiable DNA in each sample. This would control for pipetting errors, inaccurate estimates of DNA concentration, and sample-to-sample variance in the efficiency of PCR amplification (Winton et al. 2002).

However, none of these are likely to have affected the current results. For example, the low level of variability among technical replicates (Fig. 3) indicates that pipetting errors were infrequent. Variation among biological replicates was higher than for technical replicates but also was reasonably low (Figs. 4 and 6), so estimates of template DNA concentrations should have been accurate. Differences in amplification efficiency among samples were not estimated. Still, we believe they were small due to the relatively low variability among biological replicates and the smooth curves seen in the time courses. If amplification efficiencies varied among DNA samples, we would expect the points on the time-course experiments to be erratic. Instead, the points were consistent, except possibly for the 24-hour sample of Yecora Rojo. Although the results obtained from the current experiments were consistent and repeatable, including an endogenous control is an improvement that should be incorporated into future applications of this approach for estimating the quantity of *Z. tritici* DNA in infected wheat leaves.

In conclusion, this approach provided a rapid,robust, quantitative method to monitor the growth dynamics of *Z. tritici* during infection of wheat. The real-time PCR assay was more accurate and suitable for high-throughput analysis than other techniques, such as competitive PCR, which did not always correlate well with Fusarium head blight symptom expression (Nicholson et al. 1998). So far, 22 *Stb* genes for resistance to *Z. tritici* have been identified (Tidd et al. 2023). This real-time PCR assay can be used to evaluate segregating mapping populations for qualitative resistance governed by *Stb* genes and quantitative trait loci (QTL) involved in STB resistance (Adhikari et al. 2003; 2004a; 2004b; 2004c; Arraiano et al. 2001; Brading et al. 2002; Goodwin 2007; Goodwin et al. 2015; Liu et al. 2013; McCartney et al. 2002; Tidd et al. 2023; Yang et al. 2018). QTL that reduce fungal DNA could provide a more stable and long-lasting resistance in the field. Furthermore, the real-time PCR assay was used to evaluate resistance to *Puccinia coronata* f. sp. *avenae* (crown rust) in oat, which was more accurate and thorough for dissecting QTL in the Ogle/TAM O-301 (OT) mapping population compared to other assessment methods (Jackson et al. 2006; Jackson et al. 2007). More recently, regression analysis and real-time PCR assays were utilized in multi-species winter wheat field trials in Australia to assess partial resistance and to estimate genotype yield in the presence of Fusarium crown rot caused by *F. pseuodograminearum* (Milgate et al. 2023). The real-time PCR method also was valuable for detecting significant differential expression patterns of defense-related genes between resistant and susceptible cultivars and between the resistant and susceptible RILs segregating for the *Stb4* and *Stb8* genes for resistance (Adhikari et al. 2007). Overall, the real-time PCR method developed in this study is precise, specific, cost effective, and repeatable and can be a valuable tool for making decisions about managing STB disease, to identify resistant progeny in breeding programs, for investigating disease epidemiology and plant-pathogen interactions, and in assessing disease risks. Additionally, it may provide new perspectives for developing durable resistance to *Z. tritici* in wheat and thus could help reduce the threat of this pathogen to food security.

## Acknowledgements

This research was supported by USDA-ARS CRIS project 3602-22000-013-00D. We are grateful for the excellent technical assistance provided by Jill Breeden, Kristi Brikmanis, Jessica Cavaletto, Vicky Cheng, Jennifer Sanders, Ian Thompson, and S. Gurung. Judith B. Santini provided valuable guidance for statistical analyses. We also thank the researchers listed in Table 1 for providing us with either DNA or isolates of fungal pathogens tested in this research. Additionally, we express our gratitude to Lynda Ciuffetti for providing ß-tubulin gene sequences from several isolates of *Pyrenophora tritici-repentis*.

